# Next Generation of StayGold-based Adaptable Turn-On Maturation (ATOM) Sensors Targeting PSD95, Gephyrin, and HOMER1 Proteins

**DOI:** 10.1101/2025.03.17.643750

**Authors:** Harsimranjit Sekhon, Jeung-Hoi Ha, Stewart N. Loh

**Affiliations:** Department of Biochemistry and Molecular Biology, SUNY Upstate Medical University, Syracuse NY

## Abstract

Genetically encoded biosensors that have the capacity to detect intracellular molecules have proven to be invaluable tools in biology. However, development of biosensors with generalizable binding domains for detecting arbitrary targets of choice is still in its early stages. We previously introduced ATOM biosensor technology in which adaptable monobody and nanobody binding domains are employed for molecular recognition, and biosensor turn-on is achieved by coupling target binding to activation of the attached fluorescent protein by means of conformational change. In this work, we extend the ATOM mechanism to the newly identified, highly photostable fluorescent proteins mStayGold and mBaoJin. We validated the ATOM sensors using three neuronal targets: PSD95, Gephyrin, and HOMER1. The sensors exhibited turn-on ratios of over 100-fold with high specificity. Compared to existing methods for detecting these proteins, ATOM sensors demonstrated improved dynamic range, lower background, and a higher turn-on. The results highlight the adaptable nature of the ATOM mechanism and demonstrate that it allows for development of biosensors with various colors and photophysical characteristics.

## Introduction

Fluorescent protein-based biosensors (FPBs) represent some of the most powerful instruments in the neuroscientist’s toolkit. These genetically encoded sensors enable the direct visualization of molecules of interest within living cells, providing insights into their locations, concentrations, and spatiotemporal dynamics. FPBs have been instrumental in mapping neurotransmitter release^1–3^, illuminating the functions of neuronal networks^4,5^, and identifying patterns of protein expression throughout the brain^6–8^.

FPBs that transition from OFF to ON states—such as changing from dark to bright or altering colors—in response to ligand binding are particularly advantageous. The OFF state dramatically minimizes the background signal arising from free biosensors, ensuring that only sensors bound to their ligands are detectable. This feature makes it easier to identify low abundance, endogenously expressed targets in their native cellular locations.

These so-called turn-on FPBs operate through three primary mechanisms. The first, exemplified by the GCaMP family of calcium sensors (reviewed in ref^9^), involves coupling calcium ion binding to a calmodulin domain, leading to deprotonation of the chromophore in the fluorescent protein (FP) to which calmodulin and a calmodulin-binding peptide are fused. This deprotonation results in rapid and reversible activation of fluorescence. The second mechanism also requires attaching a binding domain to an FP, but it does not involve a conformational change. Instead, the activation is achieved by physically removing unbound sensor molecules from the cellular environment. This removal can occur through degradation via the ubiquitin-proteasome pathway^10–12^ or by trafficking free sensors into the nucleus, where they repress their own expression^6–8^.

We recently developed a third class of turn-on FPBs known as ‘adaptable turn-on maturation’ (ATOM) biosensors^13^ **(Fig. 1A)**. ATOM sensors incorporate a circularly permuted monobody (MB) or nanobody (NB) into one of the three surface loops or turns of an FP. Upon binding of the target ligand to the MB or NB domain, the FP chromophore undergoes maturation—spontaneous cyclization and oxidation of three consecutive amino acids—leading to an increase in fluorescence of up to 70-fold.

**Fig. 1.**
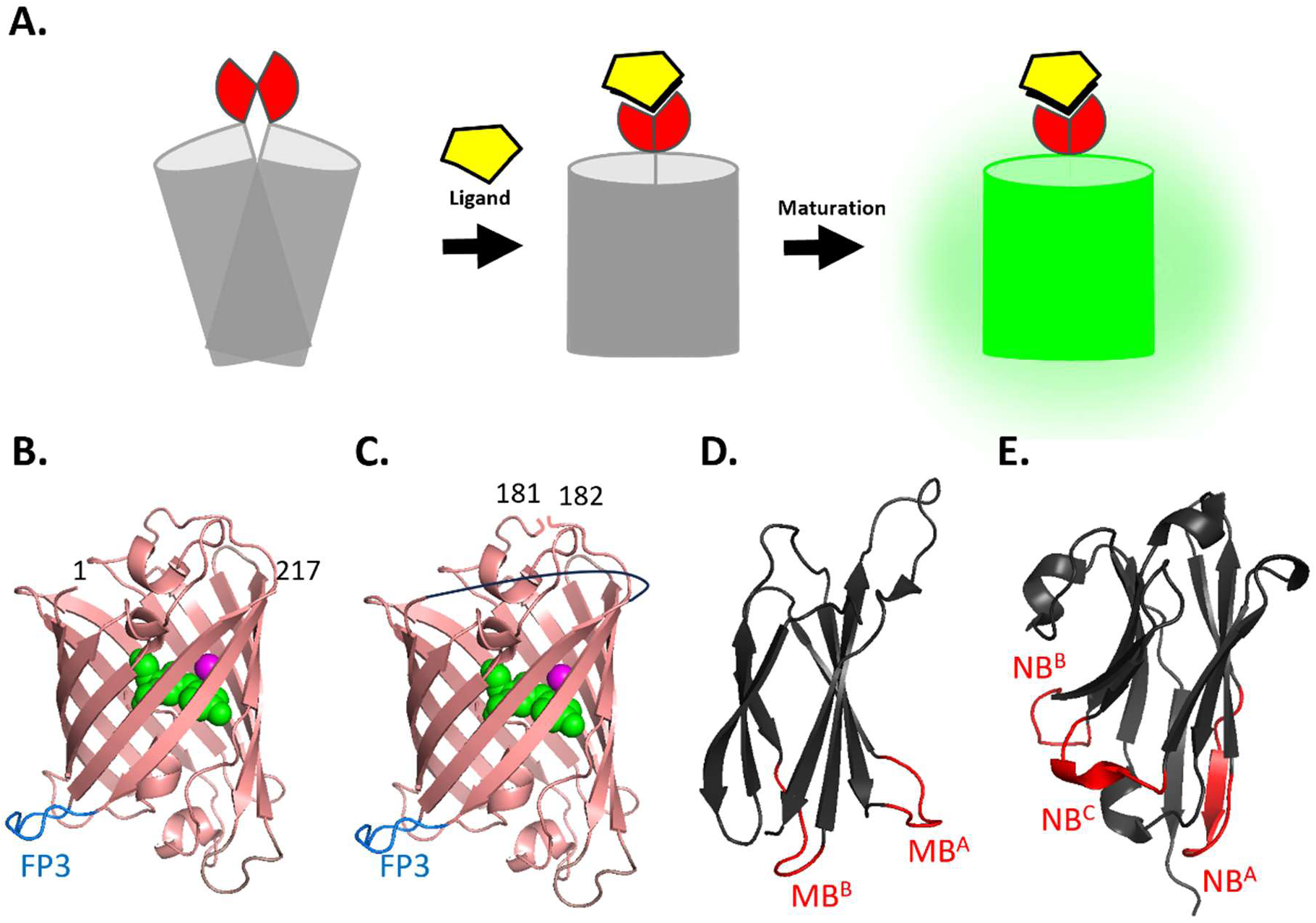
Mechanism and design of ATOM sensors. **(A)** A destabilized ligand binding domain (red) is incorporated into the fluorescent protein (FP, gray) in such a manner that the FP remains in a non-native conformation that prevents the chromophore from maturing. When the ligand (yellow) binds, it triggers the FP, allowing the chromophore to mature and activate fluorescence. Panels **(B)** and **(C)** illustrate the native (N) and circularly permuted (CP) forms of mStayGold (PDB: 8ILK), respectively, with the insertion loop FP3 highlighted in blue and chromophore shown in green. The CP structure features new N and C termini at positions 182 and 181, respectively, and a linker connecting the original N and C termini (black line). **(D)** The WDR5 monobody (PDB: 6BYN) is shown with the MB^A^ and MB^B^ loops highlighted in red. **(E)** The mCherry nanobody (PDB: 8IM0) is presented with loops NB^A^, NB^B^, and NB^C^ in red. To construct ATOM sensors, the MB or NB are circularly permuted at the loops/turns indicated in red and inserted into either the N or CP forms of the mStayGold variants.

Here, we sought to apply the ATOM design to the recently reported monomeric versions of the StayGold^14^ FP: mStayGold^15^ (mSG-J) and variants thereof, including the E138D mutant^16^ (mSG^E138D^) and the mBaoJin^17^ (mBJ) variant. Isolated initially from *Cytaeis uchidae* jellyfish, mSG-J, mSG^E138D,^ and mBJ are renowned for their improved photostability compared to other FPs. We chose to employ these biosensors to detect three proteins that are localized to the neuronal synapse and have well-established roles in synaptic function: postsynaptic density protein 95 (PSD95), gephyrin (GPHN), and HOMER1.

## Results

### Creation of SG- and BJ-based ATOM sensors and comparison with GFP-based ATOM sensors

As proof-of-concept, we first created mSG- and mBJ-based ATOM sensors that recognized WD repeat-containing protein 5 (WDR5) and mCherry (mCh) using the corresponding mClover FP-based biosensors in our earlier work as templates^13^. We inserted a circularly permuted MB (that recognizes WDR5^18^) or a circularly permuted NB (that binds mCh^19^ or the Y70A mutant that renders it nonfluorescent) into the surface loop of mSG and mBJ between strands β10 and β11 **(Fig. 1 B, C)**. We previously established two and three permutation sites for MBs **(Fig. 1D)** and NBs **(Fig. 1E)**, respectively, and three insertion sites in mClover that generated viable ATOM sensors^13^. We chose the permutation and insertion positions that had worked best (MB^A^/NB^A^ and FP3, respectively; **Fig. 1B**) in that earlier study. The mStayGold-based biosensors are designated mSGJ-ATOM^WDR5^, mSGJ-ATOM^mCh^, mSG^E138D^-ATOM^WDR5^, and mSG^E138D^-ATOM^mCh^, and the mBaoJin-based sensors are named mBJ-ATOM^WDR5^ and mBJ-ATOM^mCh^. In addition, we made a second set of biosensors that were identical except the mSG and mBJ domains were circularly permuted between strands β9 and β10, a modification that is not necessary for biosensor function but was in the original design^20^ **(Fig. 1C)**. These constructs are indicated by the prefix “CP”. **Table 1** lists all biosensors created in this study, including the binding domains and FPs from which they were constructed.

**Table 1:** Performance of ATOM biosensors for neuronal targets. Plus signs indicate turn-on with the number of (+) corresponding to the relative turn-on by eye. Negative signs indicate no apparent change in fluorescence, and double negative signs indicate turn-off. ^1^CP-mBJ-ATOM^PSD95^ ^2^mBJ-ATOM^GPHN^ ^3^CP-mBJ-ATOM^HOMER1^

| Target | Binding Domain | Reference | Addgene ID | Turn-on |  |  |  |  |  |
| --- | --- | --- | --- | --- | --- | --- | --- | --- | --- |
|  |  |  |  | CP-mBJ |  |  | mBJ |  |  |
|  |  |  |  | CP<sup>A</sup> | CP<sup>B</sup> | CP<sup>C</sup> | CP<sup>A</sup> | CP<sup>B</sup> | CP<sup>C</sup> |
| PSD95 | MB_Xph20 | Rimbault *et al.* | 133044 | ++ | ++ | N/A | + | ++ | N/A |
| PSD95 | MB_PSD95 | Gross *et al.* | 46295 | ++ | ++++<sup>1</sup> | N/A | + | ++++ | N/A |
| Gephyrin | MB_GPHN | Gross *et al.* | 46296 | ++ | - | N/A | + | ++<sup>2</sup> | N/A |
| Gephyrin | NB_GC52 | Dong *et al.* | 135224 | - | - | - | - | - | - |
| AMIGO1 | NB_AC50 | Dong *et al.* | 135225 | - | - | - | -- | -- | -- |
| HOMER1 | NB_HC20 | Dong *et al.* | 135220 | + | +++<sup>3</sup> | + | ++ | +++ | ++ |
| SAPAP2 | NB_SS80 | Dong *et al.* | 135225 | - | - | - | - | - | - |

We evaluated the new biosensors by transfecting HEK 293T cells with a plasmid encoding the biosensor gene together with a second plasmid that expressed WDR5 or the Y70A mutant of mCherry, which is recognized by the NB but is nonfluorescent **(Fig. 2)**. Turn-on ratios were calculated by measuring the total fluorescence of ATOM biosensor in cells expressing the correct ligand and dividing the mean value by that obtained from cells expressing the decoy ligand. The sensors that produced the highest turn-on ratios for WDR5 and mCh both contained the CP form of mBJ (CP-mBJ-ATOM^WDR5^ and CP-mBJ-ATOM^mCh^) (**Fig. 2A**). The fold changes were greater than or equal to those of the mClover-based ATOM sensors in our original research^13^. The second-best performers were composed of the non-permuted form of mSG-J (mSGJ-ATOM^WDR5^ and mSGJ-ATOM^mCh^).

**Fig. 2.**
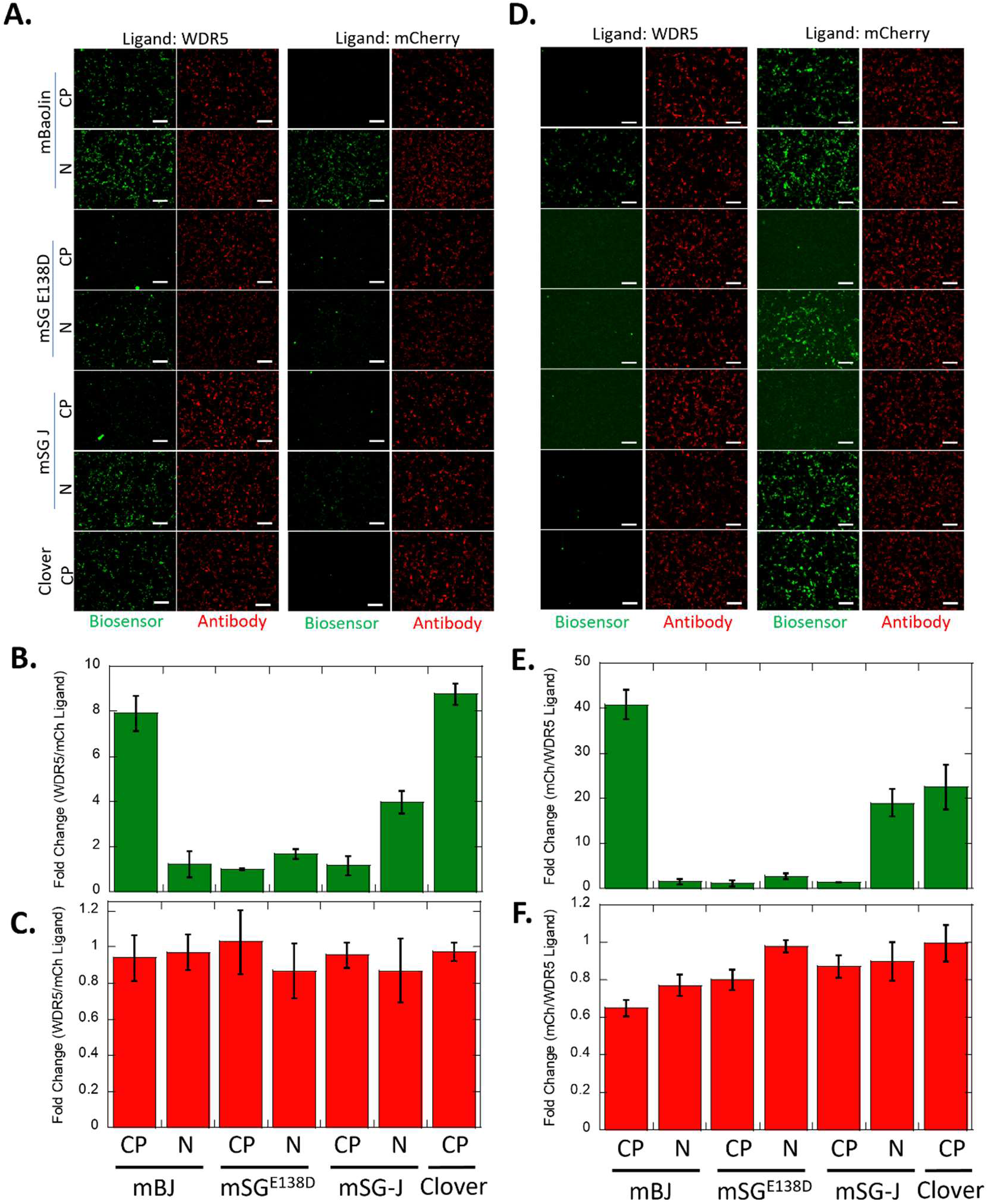
Screening for the best performing mStayGold-based ATOM biosensor. WDR5 MB^A^ **(A, B, C)** and mCherry NB^A^ **(D, E, F)** were inserted at position FP3 of either the N form or the CP form of mBaoJin, mSG^E138D^, or mSG-J. Panels **(A)** and **(D)** show representative biosensor signal (green) and total biosensor content of the cell (red) as determined by anti-HA antibody staining, in the presence of WDR5 (left two panels) or mCherry-Y70A (right two panels) ligands. The N or CP status of the FP are labeled on the left. Panels **(B)** and **(C)** show quantification of the biosensor and the antibody signal, respectively, for the WDR5 biosensors, quantified by the ratio of the biosensor or the antibody intensity in the presence of WDR5 vs. mCherry. Similarly, **(E)** and **(F)** are quantified mCherry biosensors and the antibody signals, respectively, expressed as the ratio of the intensity in the presence of mCherry vs WDR5. The data are represented as average ± standard deviation of three independent repeats, each with >100 cells. Scalebar = 100 μm.

In all instances, biosensors containing non-permuted FP domains were brighter than those with circularly permuted FP domains, though the former typically exhibited a higher background in the presence of decoy ligand **(Fig. 2A)**. Antibody staining of the biosensors revealed that they were present in cells at comparable levels irrespective of whether they were co-expressed with the correct ligand or decoy ligand **(Fig. 2C, F)**. This finding indicates that differential turnover of the sensors in their bound vs. free states did not contribute to the observed turn-on signal.

The mSG^E138D^-based WDR5 and mCherry biosensors were both dim, with the signal-to-background ratio for mSG^E138D^-ATOM^WDR5^ being so low that no turn-on was detected. We repeated the mSG^E138D^-ATOM^WDR5^ experiment using more sensitive optical-bottom multiwell dishes, which yielded a slight turn-on **(Fig. S1)**. In the original ATOM design, the overall brightness was influenced by the position of the FP into which the NB/MB was inserted, with FP1 generally being the brightest and FP3 the dimmest. We investigated whether this pattern held true for the N-SG^E138D^ protein. Surprisingly, the insertion position had minimal impact on brightness and only increased the background in the FP2 position.

To establish whether the mStayGold-based ATOM sensors had a higher photostability than our original mClover-based y-ATOM sensors, we transfected HEK 293T cells with CP-mBJ-ATOM^WDR5^ or y-ATOM^WDR5^ We then imaged the cells using a 40x objective and the highest intensity setting of the metal-halide lamp. The clover-based y-ATOM^WDR5^ sensor completely bleached within 40 seconds, whereas the CP-BJ-ATOM^WDR5^ retained >70% fluorescence after 2 minutes **(Fig. S2)**.

### Construction of PSD95, GPHN, and HOMER1 biosensors

To generate biosensors for proteins involved in cerebral cortex development, we substituted the WDR5 and mCh binding domains with MBs for PSD95^6,7^ and GPHN^6^, and NBs for GPHN, HOMER1, adhesion molecule with Ig-like domain 1 (AMIGO1), and SAP90/PSD95-associated protein 2 (SAPAP2)^21^ The nomenclature of binding domains and biosensor variants are listed in **Table 1**. The new MB and NB domains were circularly permuted at the sites shown in **Fig. 1D** and **Fig. 1E** and were inserted into position FP3 of mBJ (both non-permuted and permuted forms of mBJ; **Fig. 1B** and **Fig. 1C**). These combinations resulted in four variants for each MB-based sensor and six variants for each NB-based sensor (**Table 1**). Sensors were screened in HEK 293T cells as **in Fig. 2** with mCherry-Y70A as the decoy ligand **(Figs. S3-S7)**. The biosensors that showed the best turn-on by eye were further quantified to determine their performance.

The MBs for PSD95 and GPHN produced at least one viable biosensor. The best turn-on for PSD95 resulted from MB_PSD95 (MB^B^ permutant; **Fig. 1D**) inserted into CP-mBJ (>100x; **Fig. S8B**), and for GPHN, MB_GPHN (MB^B^ permutant) inserted in non-permuted mBJ (>5x, **Fig. S9**). These sensors are designated CP-mBJ-ATOM^PSD95^ and mBJ-ATOM^GPHN^, respectively (**Table 1**). Curiously, mBJ-ATOM^GPHN^ was >10x dimmer than CP-mBJ-ATOM^PSD95^. A possible explanation is that MB_GPHN may be less stable than MB_PSD95, which would cause GPHN_MB to thermodynamically resist the folding events that result in chromophore maturation (**Fig. 1A**). Another possible reason may be low overall cellular expression of the GPHN ligand in HEK 293T cells.

Compared to the MB-based biosensors, those made from NBs had a lower success rate. In most cases, they were either always bright or always dim, regardless of ligand. The bright group generally consisted of NBs inserted into non-permuted mBJ whereas in the dim group the NBs were inserted into CP-mBJ. Interestingly, for AMIGO1 we observed an always dim phenotype when the NBs were inserted in CP-mBJ, but a turn-OFF phenotype when they were inserted in mBJ **(Fig. S5)**. Although this result is not readily explainable, it is not unprecedented, as we reported a similar phenomenon with the hRas-binding MB in our original study^13^. The only NB-based sensors that produced high turn-on ratios employed the HOMER1 NB (NB_HC20) fused to either mBJ or CP-mBJ **(Fig. S6)**. Of these, the NB^B^ permutant inserted into CP-mBJ (designated CP-mBJ-ATOM^HOMER1^; Table 1) performed best (>50x turn-on; **Fig. S10**). In summary, we successfully created ATOM biosensors for PSD95, GPHN, and HOMER1, but failed to generate sensors for AMIGO1 and SAPAP2.

### Comparing ATOM sensors to FingR sensors

Next, we sought to compare specificity and response of the PSD95, GPHN, and HOMER1 ATOM sensors to that of the well-established FingR biosensors. Briefly, the FingR sensors contain a transcriptional repressor domain fused to the MB recognition domain such that any unbound sensor translocates to the nucleus and inhibits its own transcription^6^. We transfected HEK 293T cells with a plasmid encoding PSD95, GPHN, or HOMER1 and a second plasmid that either expressed one of the three corresponding ATOM biosensors or the published FingR biosensor for PSD95 or GPHN. Measuring the average total fluorescence of the cells in each well provided a test of specificity of the biosensors as well as a quantitative comparison of ATOM and FingR performance.

All three ATOM sensors produced a high turn-on (450 ± 70 for PSD95, 14.1 ± 3.3 for GPHN, and 90 ± 15 for HOMER1; **Fig, 3, left**), whereas the FingR biosensors responded more modestly (2.7 ± 0.6 for PSD95 and 2.6 ± 0.4 for GPHN; **Fig. 3, right**). The lower turn-on of FingR sensors was due to high nuclear background fluorescence, which contained most of the unbound sensor. We note, however, that the FingR sensors have a very low cytoplasmic background, so the turn-on ratio is much higher in images where the nucleus can be excluded. For example, neuronal processes project far from the nucleus, and the FingR exhibited 50x turn-on for detecting PSD95 in dendrites and synapses of mouse cortical neurons^6^.

**Fig. 3.**
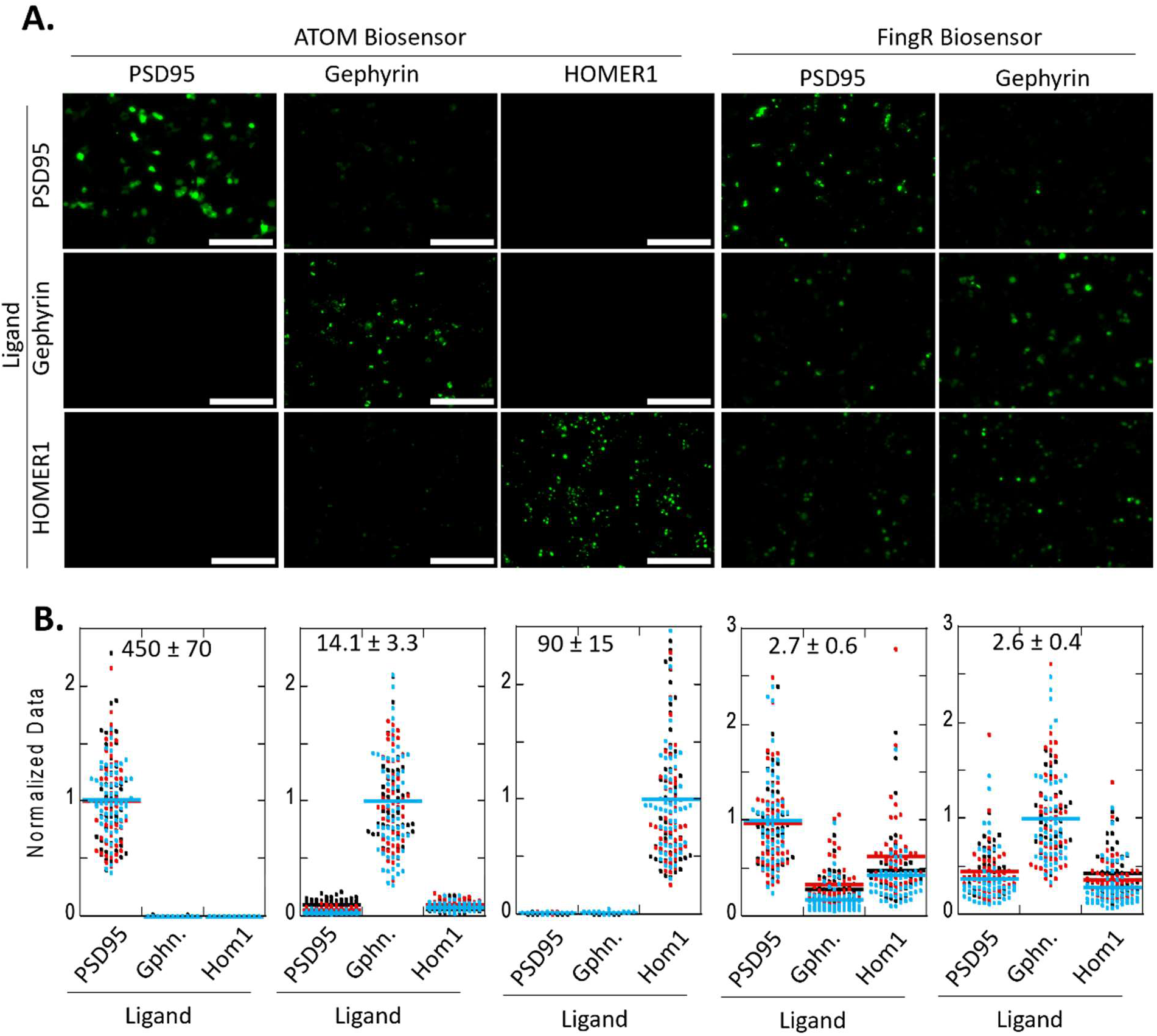
Development of ATOM sensors for neuronal targets. **(A)** Raw images of ATOM and FingR biosensors co-transfected with either PSD95, GPHN, or HOMER1 ligands. The biosensor is labeled on the top and the ligand on the left. **(B)** Quantification of the biosensors above. Each dot represents a cell, and the different colors represent three independent repeats. The average fold turn-on and the standard deviation of three repeats are provided. Scalebar = 100 μm.

We next compared the linearity of responses of the ATOM and FingR sensors by co-transfecting varying amounts of DNA encoding the PSD95 ligand together with a fixed amount (400 ng) of plasmid DNA expressing either CP-mBJ-ATOM^PSD95^ or the FingR PSD95 biosensor **(Fig. 4A)**. The ATOM signal was proportional to PSD95 plasmid concentration over the range of 2 ng to 150 ng **(Fig. 4B)**. By contrast, the response of FingR was nearly flat over the same concentration range, showing minimal activation. We reasoned that result was likely due to the high background fluorescence coming from unbound FingR sensor in the nucleus.

**Fig. 4.**
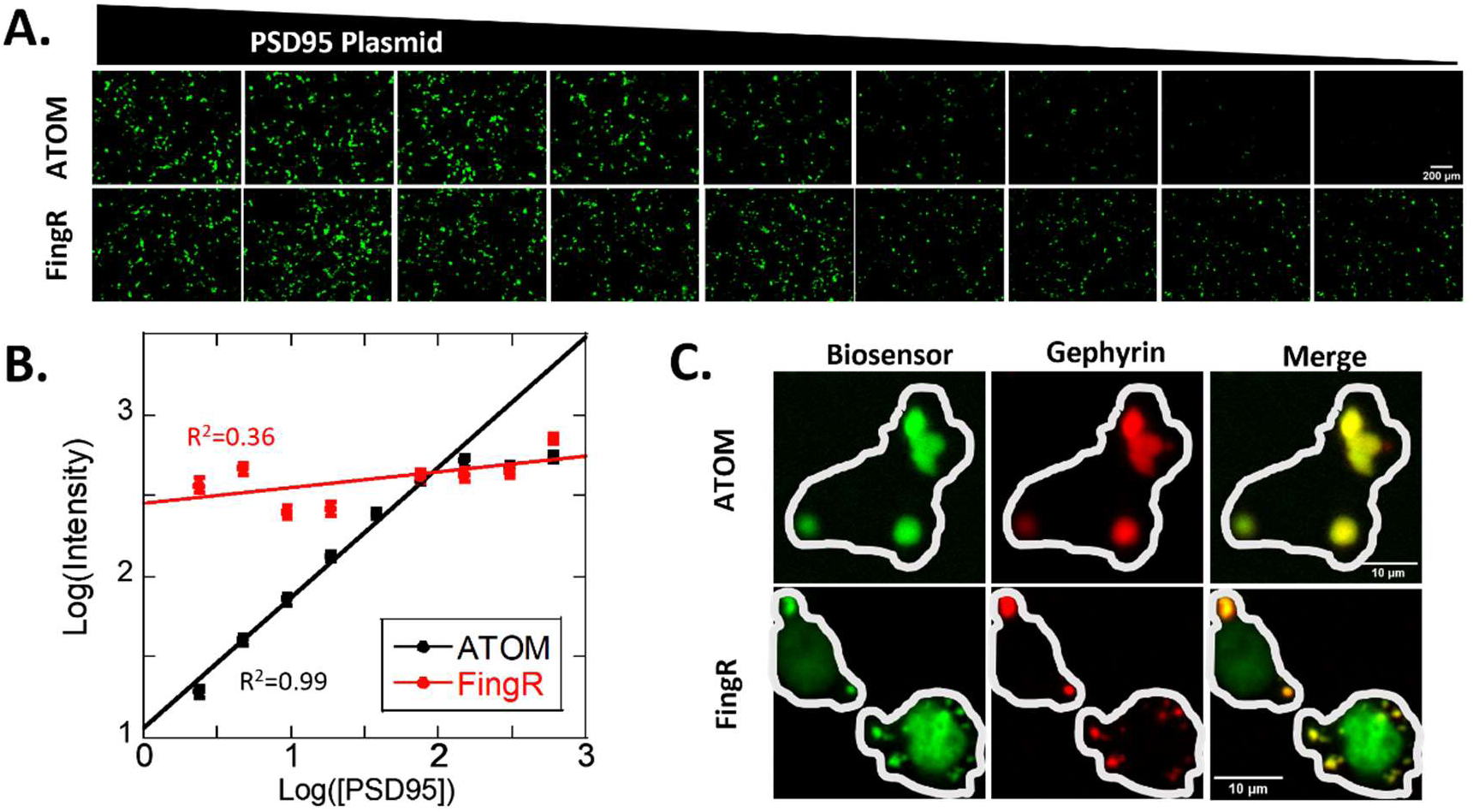
Comparison of ATOM and FingR. **(A)** Raw representative images of a fixed amount (400 ng) of plasmid DNA expressing either the ATOM (top) or FingR (bottom) biosensor co-transfected with various amounts of plasmid DNA encoding the PSD95 ligand (600 ng on the left, diluted 2x from the previous amount for all the images on the right). **(B)** Each data point represents average ± standard error of the experiments above, plotted as a function of PSD95 plasmid concentration. A linear fit was performed, excluding the highest two PSD95 plasmid concentration points as those data were determined to be saturating by eye. The R^2^ values are shown for each fit. ATOM had a linear response to PSD95 plasmid concentration whereas the FingR biosensor changed minimally. **(C)** representative images of the GPHN ATOM (top) and FingR (bottom) biosensors in green, and GPHN ligand fused to miRFP647 in red. The white cell outline was drawn manually after enhancing the contrast of the images. ATOM co-localizes to the GPHN ligand whereas FingR shows a high nuclear background fluorescence. The experiments are representative of two biological repeats.

To verify the above conclusion, we took advantage of the observation that GPHN expressed as extranuclear puncta whereas PSD95 expressed evenly throughout in the cytoplasm in HEK 293T cells. We co-transfected HEK 293T cells with three plasmids encoding the GPHN ligand (tagged with miRFP670 for visualization), and two additional plasmids expressing mBJ-ATOM^GPHN^ and the corresponding FingR biosensor. miRFP670 fluorescence confirmed that GPHN was located exclusively in cytoplasmic puncta and not in nuclei, as expected (**Fig. 4C**). FingR fluorescence overlapped with the puncta but also localized strongly to nuclei. ATOM fluorescence also coincided with the puncta but was at background levels in nuclei, exactly matching the pattern of miRFP670 fluorescence. We conclude that ATOM removes the nuclear fluorescence artifact of FingR, allowing for unambiguous determination of cellular locale and better visualization of targets that are near the nucleus.

Curiously, we consistently observed that the PSD95 FingR sensor influenced the localization of the PSD95 ligand in HEK 293T cells, a phenomenon not seen with CP-mBJ-ATOM^PSD95^. When cells were co-transfected with plasmids encoding the FingR sensor and the PSD95 ligand, PSD95 exhibited a more punctate distribution compared to the control (cells transfected only with the plasmid encoding PSD95 ligand), in which PSD95 showed diffuse cytoplasmic expression. Some puncta were even extracellular, suggesting PSD95 was being extruded from the cell (**Fig. S12**). In contrast, PSD95 retained its diffuse expression pattern in cells co-transfected with the CP-mBJ-ATOM^PSD95^ plasmid.

### Determining the turn-on mechanism of StayGold-based ATOM sensors

To establish whether degradation of unbound biosensors contributed to the turn-on mechanism of the StayGold-based ATOM biosensors, we fused a 2x-HA tag to the sensors to report on total cellular expression. HEK 293T cells were transfected with a plasmid encoding the mSG, mSG^E138D^, and mBJ ATOM sensors for WDR5 and mCherry and a second plasmid expressing either WDR5 or mCherry. HA antibody staining indicated that expression of all mStayGold-based ATOM sensors remained the same regardless of whether the sensors were co-expressed with the correct or decoy ligands. **(Fig. 2)**. Additionally, we repeated the titration experiment in which HEK 293T cells were co-transfected with a fixed amount of plasmid encoding CP-mBJ-ATOM^PSD95^ and a variable amount of plasmid expressing PSD95, but this time stained with HA antibody. Notably, while biosensor fluorescence correlated well with the expression of PSD95 ligand, the antibody signal remained constant and was independent of ligand concentration. Hence, the activation mechanism of StayGold-based ATOM sensors, like that of their mClover-based counterparts, appears to involve ligand-induced conformational changes (**Fig. 1A**) as opposed to reduction of biosensor turnover.

## Discussion

Our previous research introduced ATOM, a mechanism for the maturation of fluorescent protein chromophores that can be integrated with the molecular recognition of binding domains to create a biosensor whereby ligand binding activates fluorescence^13^. In this study, we expand upon that mechanism by exploring a newly discovered, highly photostable lineage of fluorescent proteins derived from StayGold. After comparing the three recently reported monomeric variants of StayGold, we identified mBaoJin as the most effective for constructing ATOM biosensors. Building on these insights, we developed sensors targeting neuronal proteins PSD95, GPHN, and HOMER1.

ATOM offers several distinct advantages over other methods for detecting intracellular ligands that utilize generalized binding domains. Compared to the widely used FingR class of biosensors, ATOM not only achieves a higher turn-on ratio due to lower nuclear background (when total cellular fluorescence is integrated),but may also reduce the likelihood of the biosensor perturbing ligand localization. Verkhusha and colleagues recently described an alternative mechanism in which the biosensors are inherently unstable in the absence of target, leading to their degradation via the ubiquitin-proteasome system^10,11^. Turn-on is achieved by suppression of this default “turn-off” activity by reduced turnover in the presence of ligand. This mechanism was also previously employed by Cepko and co-workers^12^, with the primary distinction being how the NB domains were made unstable (the Verkhusha approach involved inserting a far-red FP within the nanobody, while the Cepko method introduced destabilizing mutations within the nanobody). Unlike these turnover-dependent biosensors, ATOMs do not degrade in the absence of their targets, making them suitable for use in cellular compartments that lack degradation machinery. We note that both ATOM and turnover-type biosensors involve manipulations to the binding domain such as circular permutation, insertion of an FP, or mutations to enhance instability, which may lead to reduced binding affinity. Reduced affinity is less of a concern with FingR sensors, due to their straightforward design that fuses the wild-type binding domain with a transcriptional repressor.

One drawback of ATOM sensors compared to FingR and turnover-type biosensors is that, because the ATOM conformational change mechanism is poorly understood, it’s necessary to screen a 6-9 member library of MB/NB permutation sites and FP insertion points. Future studies will aim to elucidate the ATOM mechanism in greater detail, facilitating the rational design and construction of ATOM biosensors.

We previously demonstrated that ATOM is compatible with FPs originating from *A. victoria* jellyfish and *E. quadricolor* sea anemone, and we have now incorporated a third lineage into our toolkit. This finding highlights a conserved mechanism among multiple FP ancestries that enables construction of ATOM biosensors. A recent paper outlines protocols for constructing ATOM sensors from FPs and MBs, NBs, and potentially other types of binding domains^22^. These protocol steps were followed to develop the current StayGold-ATOM sensors. We recommend that readers consult those instructions to exploit the ATOM design to create sensors for novel targets or new FPs.

## Methods

### Gene Construction

The WDR5 ATOM biosensors were cloned into the pCAG vector containing a CAG promoter along with the MBP tag in the following sequence: MBP—2xHA—ATOM. An EcoRI site was inserted at the 5’-end of the MBP, followed by an AgeI site between MBP and the HA tags. A NotI site was inserted at the 3’-end of the ATOM gene. To clone the different mStayGold variants, we used overlapping extension PCR to insert the CP MB at the FP3 position and inserted the vector using the AgeI and NotI sites. The mCherry ATOM sensors were cloned in the same way except the pCMV-N1(Clontech) vector was used, containing the CMV promoter.

The constructs used for screening the neuronal biosensors were directly cloned into the pCAG vector without an HA tag or MBP, using overlapping PCR and the EcoRI and NotI restriction sites. The best-performing biosensors were then moved to the same pCAG-MBP-2xHA vector as described above.

The parental nanobodies and nanobodies were obtained from the following Addgene submissions: PSD95 MB 1 (Addgene 133044, gift from Matthieu Sanlos); PSD95 MB2 (Addgene 46295) and GPHN MB (Addgene 46296) were both gifts from Don Arnold; GPHN NB (Addgene 135224), SAPAP 2 NB (Addgene 135288), HOMER1 NB (Addgene 135220), and AMIGO1 NB (Addgene 135225) were all gifts from James Trimmer. The ligand plasmids were obtained from the following add gene submissions: GPHN1 (Addgene 68820, gift from Shiva Tyagarajan), SAPAP2 (Addgene 40216, gift from Yutaka Hata and Yoshimi Takai), HOMER1c (Addgene 89442, gift from Matthew Kennedy), and AMIGO1 (Addgene 157350, gift from Chris Garcia). The mStayGold and mBaoJin genes were synthesized by Eurofins Genomics. The WDR5 and mCherry-Y70A genes were cloned as previously described. The PSD95 gene was a gift from Dr. Eric Olson.

The PSD95 gene was cloned both alone and fused to bright mCherry for localization in the pCAG vector as described above. The HOMER1c gene was fused to miRFP670 using the NheI and SacI sites in the parental vector. The GPHN gene was also fused to miRFP670 using the BsrGI and NheI sites in the parental vector. The AMIGO gene was fused to miRFP670 using NheI and SacII sites downstream of the miRFP670 in the pCAG vector.

### Cell Culture and transfection

HEK293T (CRL-3216) cells were purchased from ATCC and cultured at 37°C in DMEM (with GlutaMax) containing 10% FBS and 1x penicillin-streptomycin. The cells were split every two days at a 1:5 dilution and discarded after passage 20. For screening in figs. 1, S3-S7, the cells were split onto a plastic 24-well dish (Nest catalog number 702001). All other experiments utilized a glass-bottom 24-well dish (CellVis catalog number P24-1.5H-N). Briefly, the cells were split 1 day before transfection such that the confluency at the time of transfection was ∼80%. Transfection was performed the following day using calcium phosphate as previously described^13,20,23^, using 500 ng total DNA unless otherwise described in the figure captions. In cases where multiple plasmids were transfected, they were combined in equimolar ratios, and 500 ng of the mix was transfected. The cells were either imaged in imaging media (DMEM, no FBS) or fixed for antibody staining 36-48 hrs after transfection.

### Antibody Staining

We coated the plates with collagen before plating the cells for all the experiments requiring antibody staining. A solution of 1:1000 dilution of glacial acetic acid and 20 μg/mL collagen was prepared in autoclaved ddH_2_O, and 1 mL of this solution was dispensed into each well of the 24-well plate. After 8 hr-overnight incubation at 37°C, the solution was discarded, and the plates were rinsed with autoclaved ddH_2_O three times. The plates were then dried in the hood for at least 2 hr and sterilized using the UV lamp for 30 min.

After 36-48 hr of transfection, the cells were briefly rinsed with PBS, followed by a 30 min fixation in 3 % paraformaldehyde while shaking. Three wash steps, 5 min each, were then performed using 20 mM Tris pH 7.5, 150 mM NaCl. The cells were then permeabilized and blocked in the blocking buffer (5% BSA, 0.25% Triton X-100, 20 mM Tris pH 7.5, 150 mM NaCl) for 30 min. The anti-HA primary antibody (XX) was diluted 1:1500 in the blocking buffer and the cells were labeled for 1 hr while shaking, followed by three washes of 20 mM Tris pH 7.5, 150 mM NaCl. The secondary antibody conjugated with Alexa-594 was also diluted 1:1500 in blocking buffer and the cells were labeled for 30 min, followed by three 5 min washes in PBS. The plates were then either imaged promptly or stored at 4°C overnight.

### Imaging and analysis

The images were acquired using a Zeiss Axiovert 200 m microscope. A 10x/0.3 Plan-NEOFLUAR objective was used for images in figs 2, 4A, S1, S3-S11, 20x/0.4 ACHROPLAN for fig. 3, 40x/0.75 Plan-NEOFLUAR Ph2 for figs S2, S12, and 100x/1.4 Plan-APOCHROMAT DIC for fig. 4C. The excitation source was a metal-halide lamp. A 500/20 excitation, 515 Dichroic, 535/15 emission filter was used to acquire biosensor images and 546/12 excitation, 580 dichroic, 590/high pass filter was used to acquire the antibody staining images. The lamp intensity and integration time were determined using the positive controls and kept the same for all the conditions for a specific biosensor.

For each experiment, images were captured from a minimum of three fields of view, with each field containing at least 30 cells, and all three images were combined for quantitative analysis. The images were processed using Fiji^24^ software. In cases where there was no accompanying antibody staining channel, cell intensities were quantified manually by drawing a region of interest around each cell and integrating the intensity. When a reference antibody staining channel was available, cells were first segmented using this channel with a custom ImageJ macro that utilized Otsu’s thresholding method. The intensities in both the biosensor and antibody channels were then measured based on the segmented cells. Prior to quantification, both methods required background correction, which was performed using ImageJ’s built-in plugin with a 50-pixel sliding ball radius.

### Reproducibility and Data Availability

All experiments described in this study were performed at least two times, with the exact number provided in figure legends. For biological repeats, the experiments were conducted on different days with cells freshly split and transfected each time. All the raw data collected in this study will be provided on a FigShare repository upon the acceptance of this manuscript. The plasmids generated here will be provided upon request until deposited on Addgene.

## Supplementary Images

**Fig. S1.**
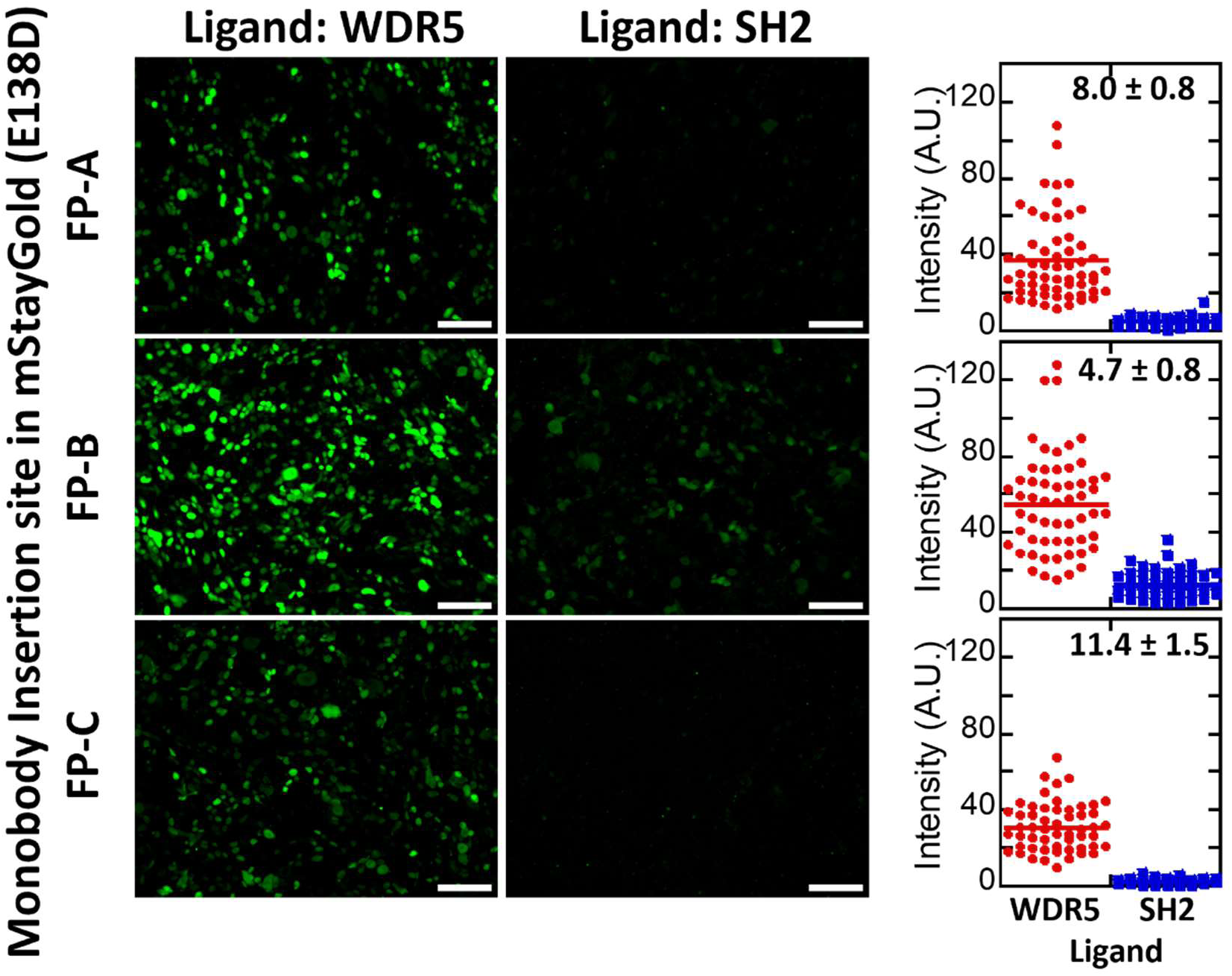
WDR5 monobody inserted in other loops of mSG^E138D^. To enhance the brightness and activation of the mSG^E138D^ biosensors, we incorporated the WDR5 MB1 at all the positions evaluated in our original study with mClover. However, the resulting biosensors did not exhibit significant improvement in brightness compared to the original FP-C insertion position and, in fact, performed worse. SH2 was chosen as the decoy ligand. The quantification on the right is representative of three biological repeats with the average ± standard deviation of the three repeats stated on the top right. Scalebar = 100 μm.

**Fig. S2.**
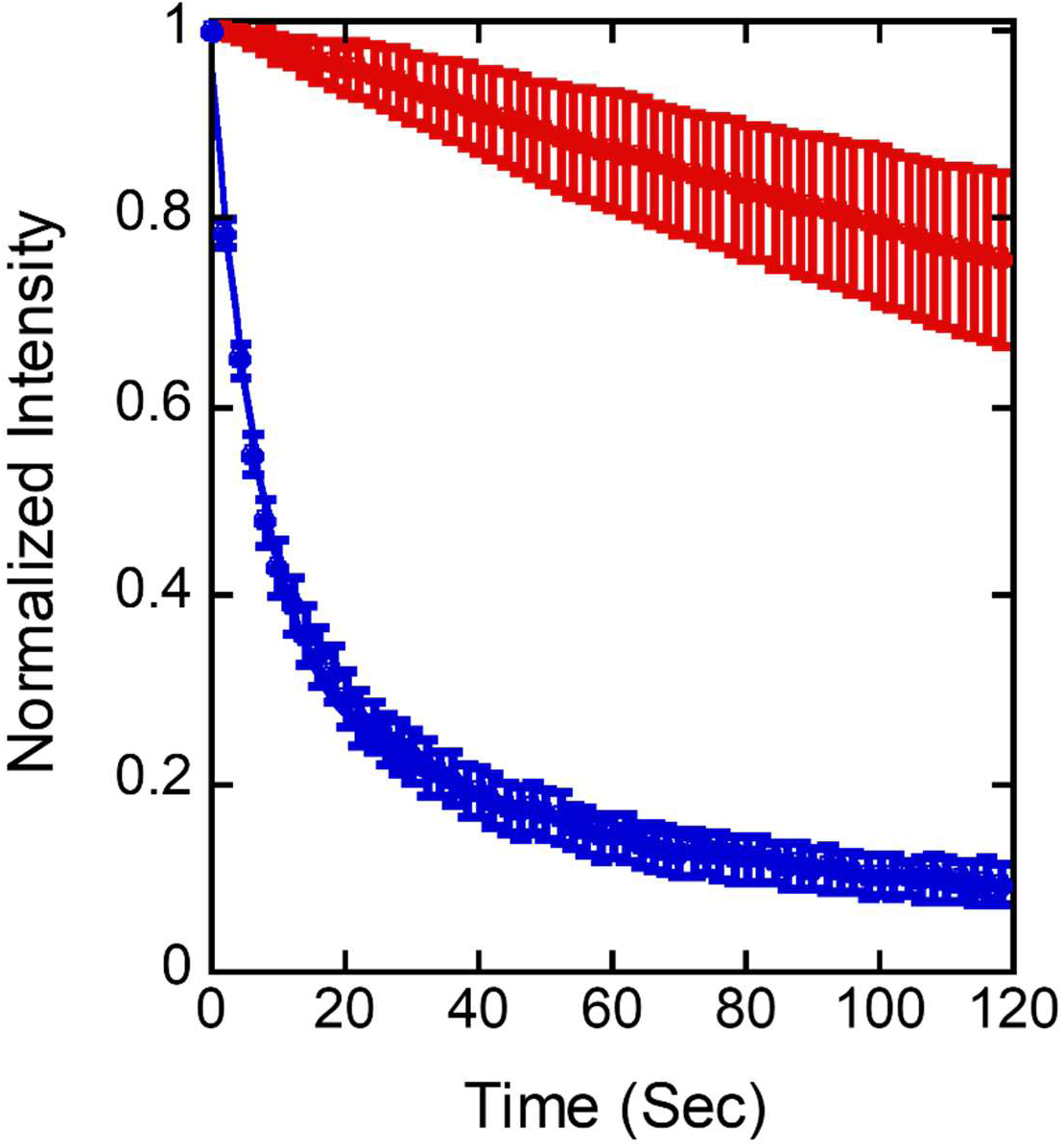
Photostability of StayGold derived ATOM biosensors. CP-mBJ-MB^WDR5^ (red) and CP-mClover-MB^WDR5^ (blue) biosensors co-transfected with the WDR5 ligand and kept under the highest lamp intensity. The mClover-based biosensor bleaches completely within 2 minutes, whereas the mBJ-based sensor retains ∼80% of its fluorescence. Fits are an exponential decay + linear line.

**Fig. S3:**
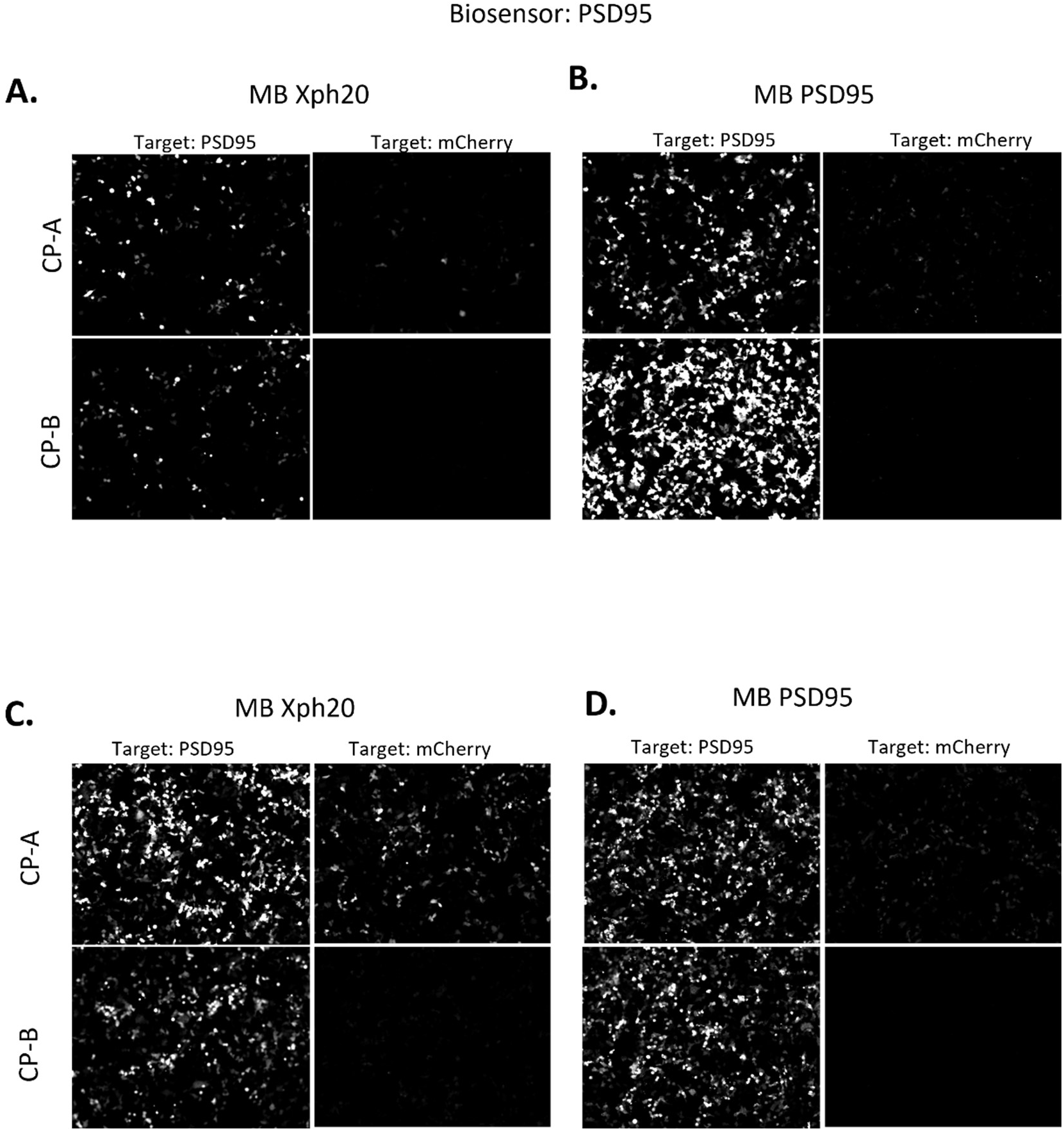
Screening the PSD95 Biosensors. For both the monobodies for PSD95 (Monobody 1-**A, C** and 2-**B, D**), both CP-A and CP-B were inserted in either the CP frame **(A, B)** or the N frame **(C, D)** of mBaoJin.

**Fig. S4:**
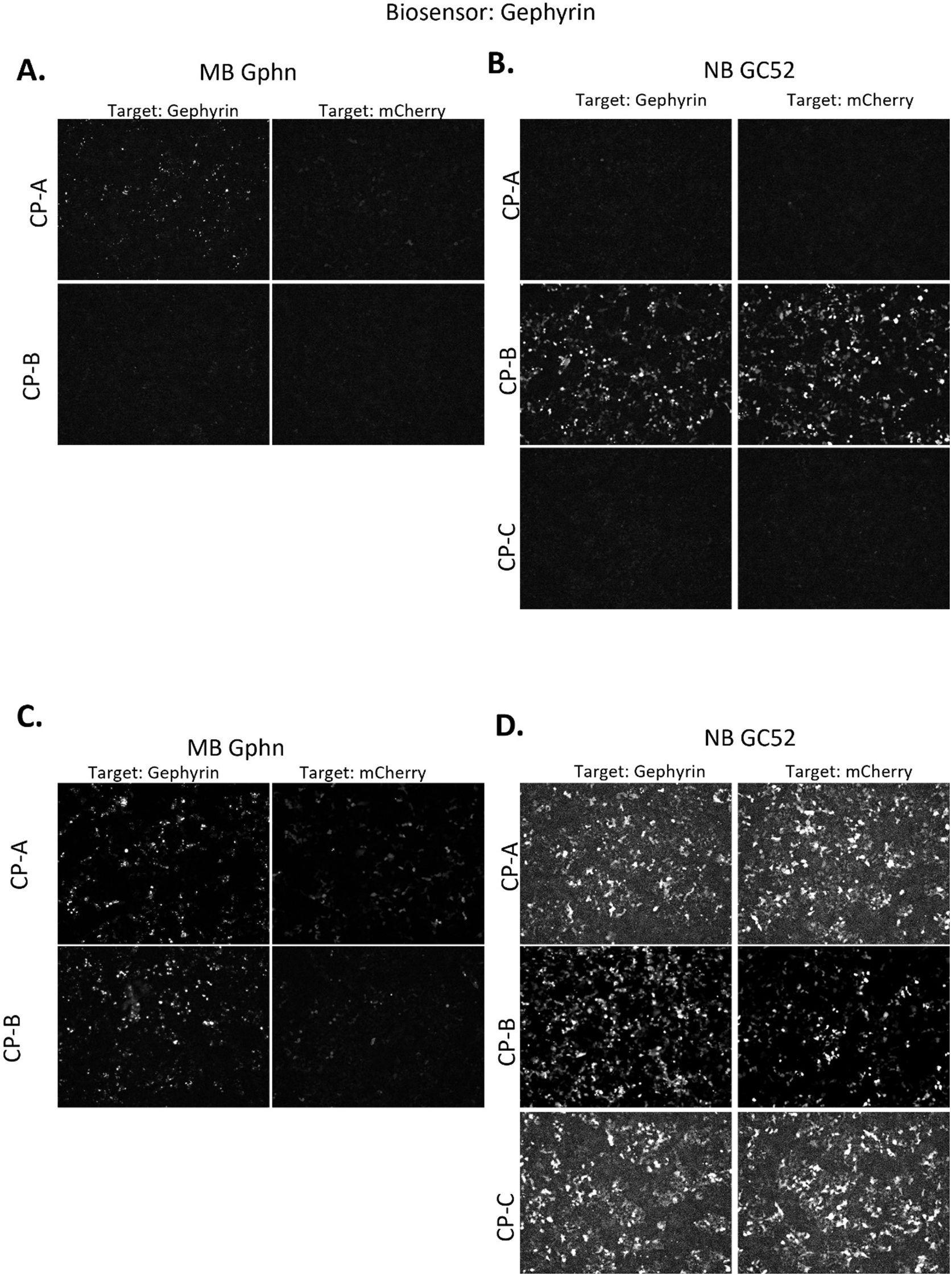
Screening the GPHN Biosensors. All the CP positions for the GPHN monobody **(A, C)** or nanobody **(B, D)** described in the text inserted in either the CP frame **(A, B)** or the N frame **(C,D)** of mBaoJin.

**Fig. S5:**
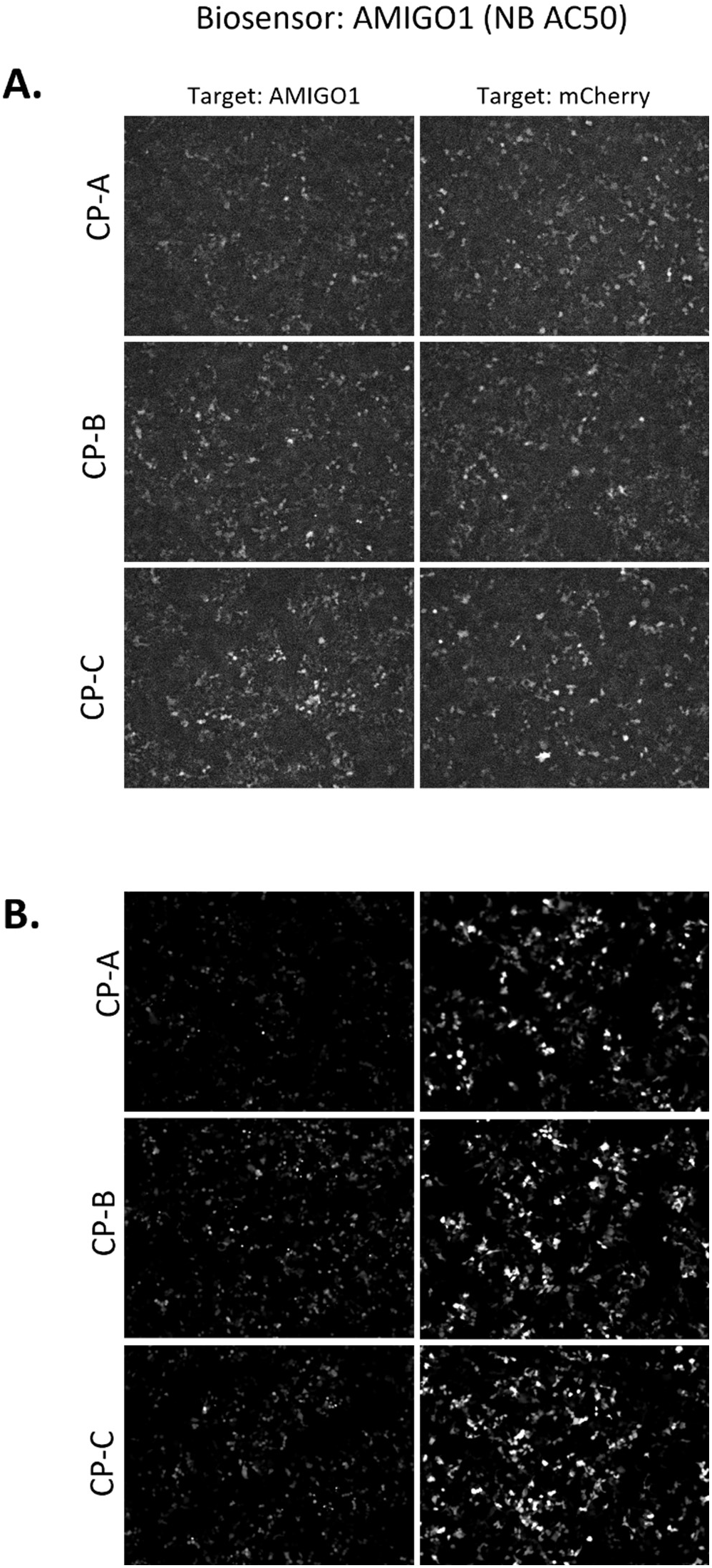
Screening the AMIGO-1 Biosensors. The three CP positions for the AMIGO-1 nanobody inserted in either the CP frame **(A)** or the N frame **(B)** of mBaoJin.

**Fig. S6:**
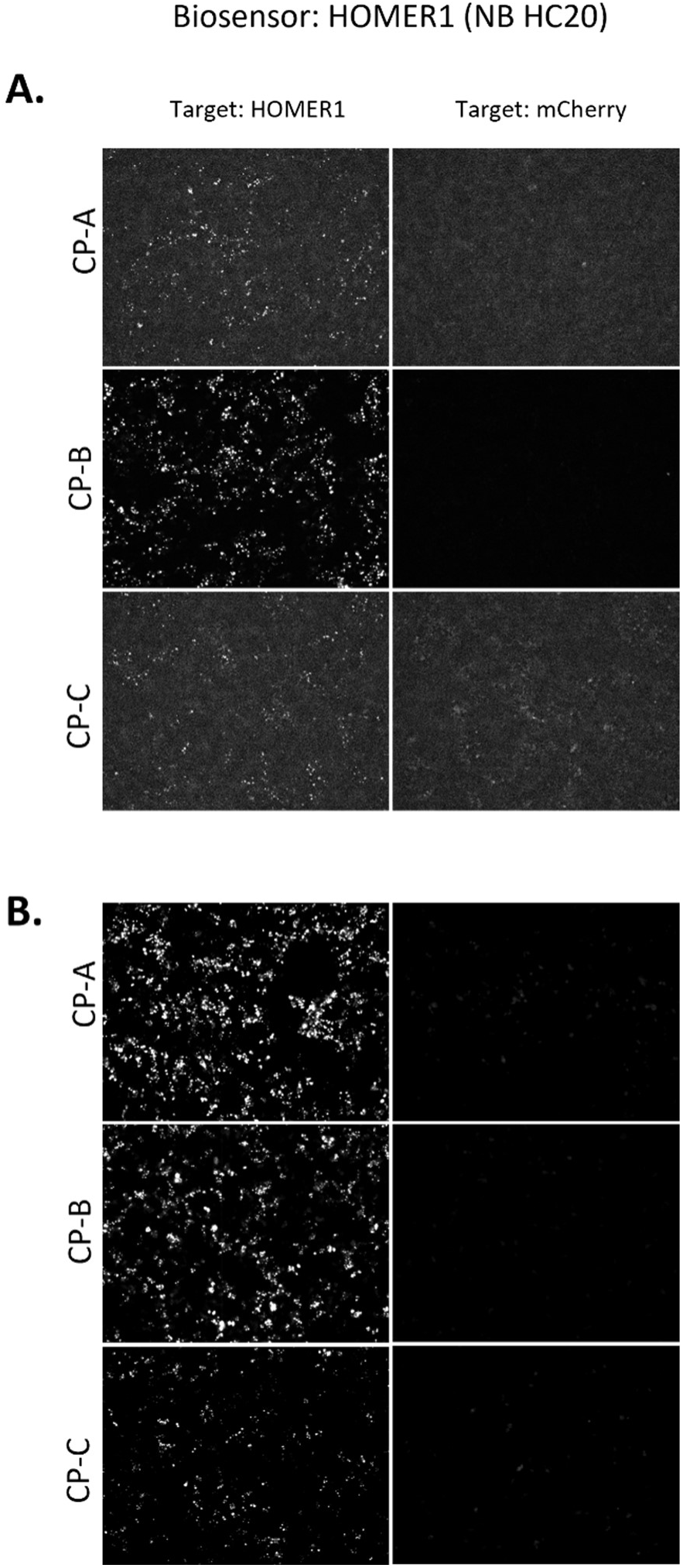
Screening the HOMER1 Biosensors. The three CP positions for the HOMER1 nanobody inserted in either the CP frame **(A)** or the N frame **(B)** of mBaoJin.

**Fig. S7:**
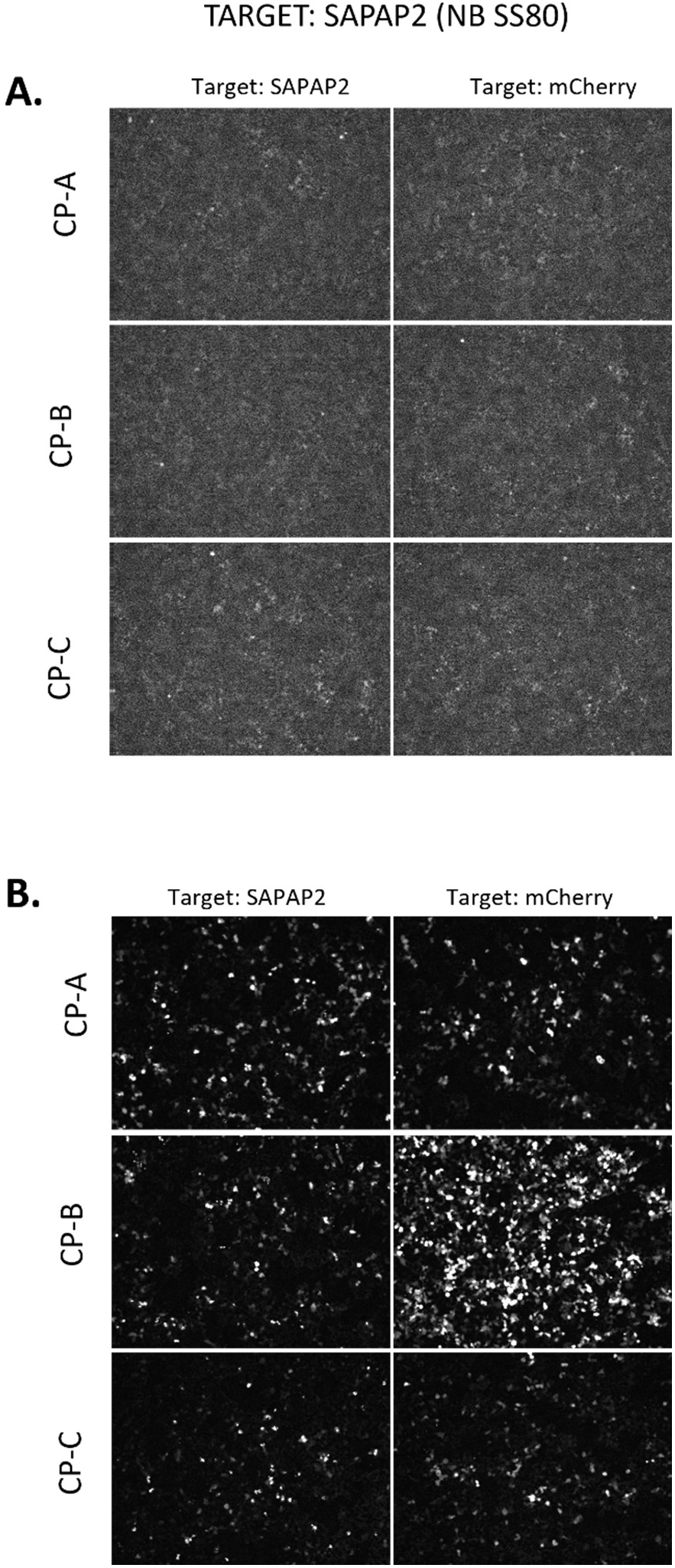
Screening the SAPAP2 Biosensors. The three CP positions for the SAPAP2 nanobody inserted in either the CP frame **(A)** or the N frame **(B)** of mBaoJin.

**Fig. S8:**
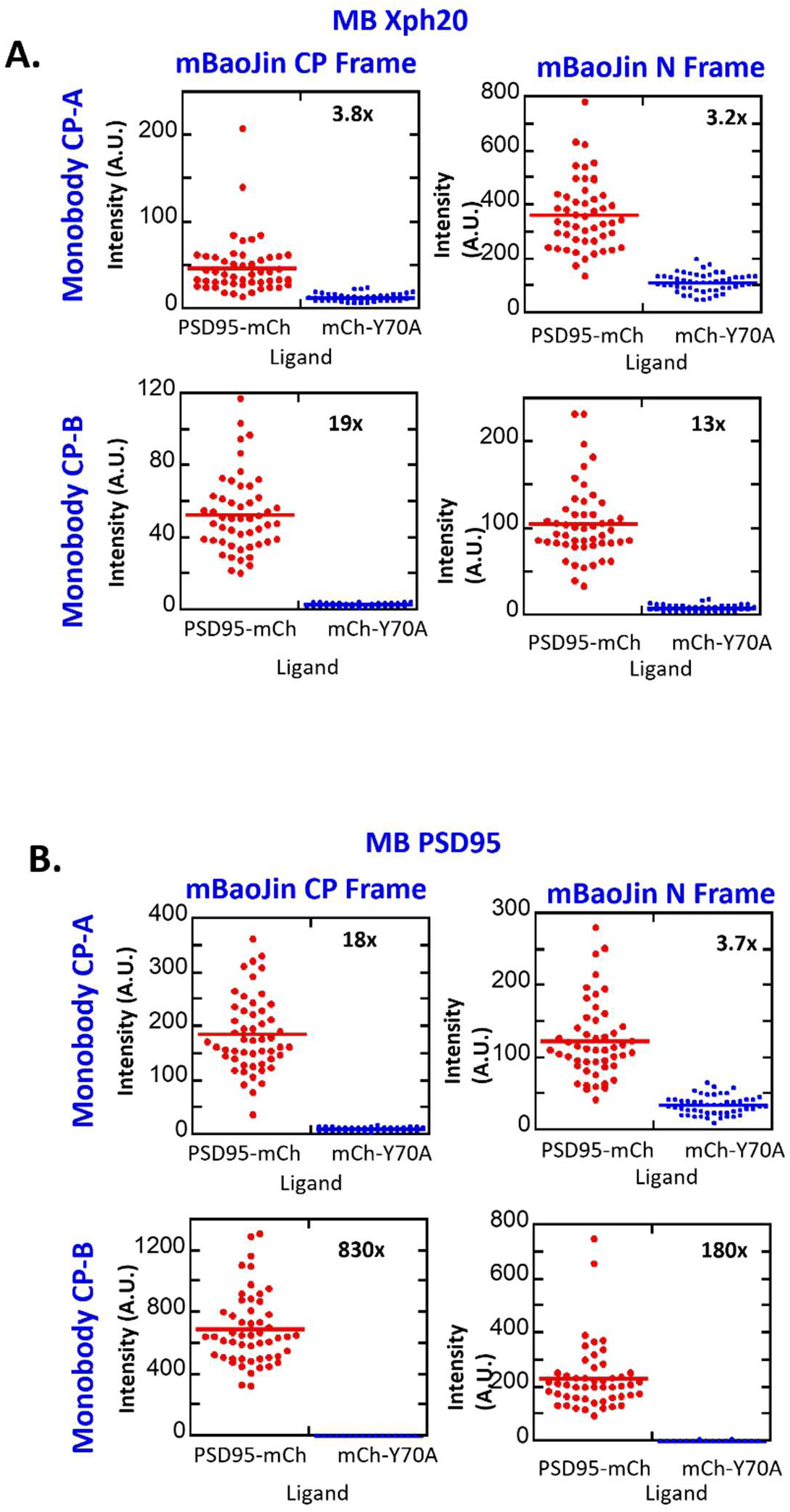
Quantification of PSD95 Biosensors. The PSD95 biosensors made from Monobody 1 **(A)** or Monobody 2 **(B)** from fig. S3 quantified. The CP or the N frame of mBaoJin is labeled at the top, and the CP position of each monobody is labeled on the left. In both cases, the CP-B monobodies produced the highest turn-on, with Monobody 2 outperforming Monobody 1.

**Fig. S9.**
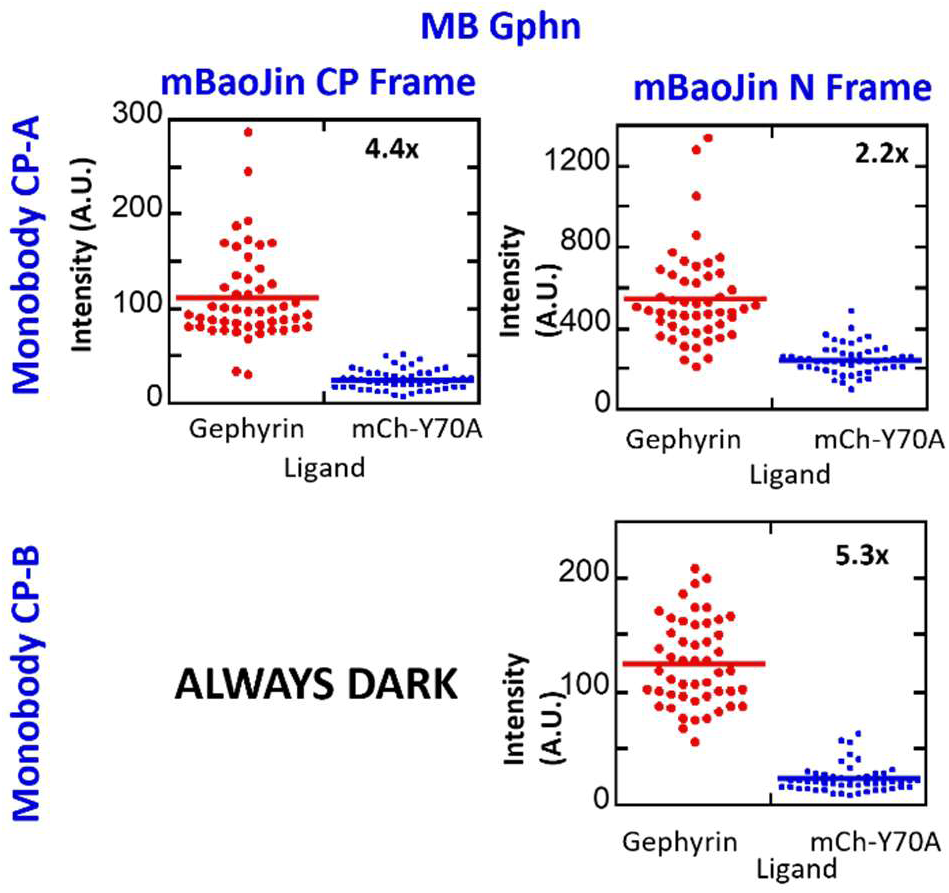
Quantification of GPHN Biosensors. The GPHN biosensors made from the monobody in Fig. S4 quantified, with the same labeling scheme as Fig. S8. The CP-B Monobody, when inserted in the CP frame of mBaoJin, was always dark, whereas the same monobody, when inserted in the N frame, slightly outperformed CP-A.

**Fig. S10.**
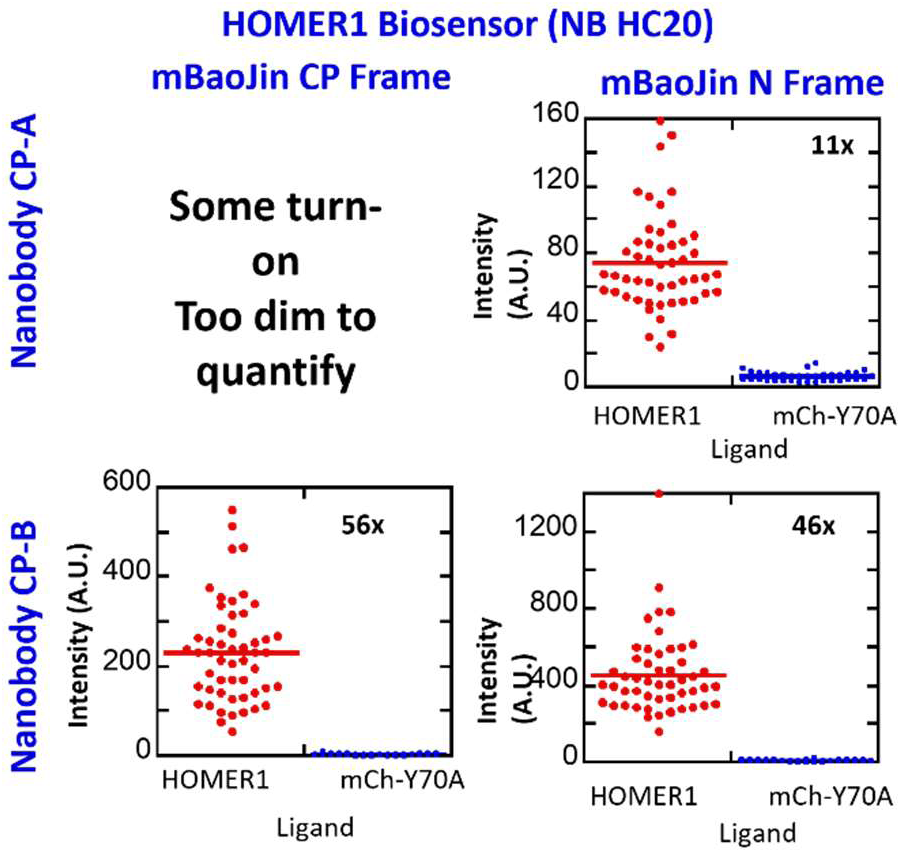
Quantification of the HOMER1 Biosensors. The HOMER1 biosensors made from nanobody from Fig. S6 quantified, with the same labeling scheme as Fig. S7. The CP-C nanobody was deemed too dim by the eye and was, therefore, not quantified. CP-B Nanobody in both the CP and the N frames of mBaoJin had the highest turn-ON.

**Fig. S11.**
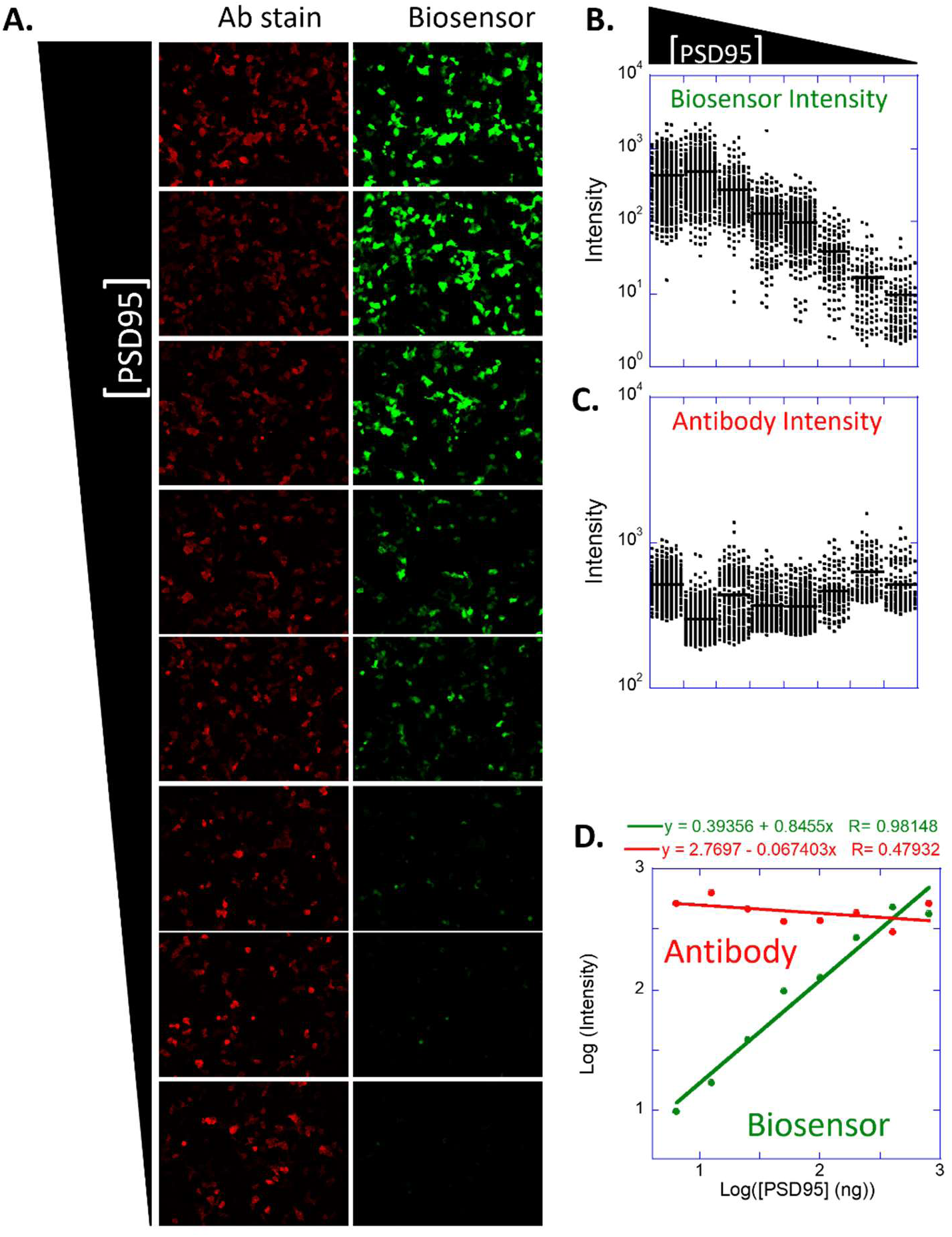
The same experiment in Fig. S4 was repeated along with antibody staining for the HA tag present in the biosensor. **(A)** representative raw images of antibody stain biosensor fluorescence when co-transfected with 400 ng of the biosensor and titration of the PSD95 ligand, 600 ng at the top image, diluted 2x from the previous image going down. **(B)** and **(C)** shows quantification of the biosensor intensity and the antibody intensity, respectively. Each dot represents a cell. **(D)** Antibody or biosensor intensity plotted as mean ± standard error of the above data as a function of PSD95 concentration. The biosensor intensity increases linearly with rising ligand concentration, while the antibody fluorescence remains constant.

**Fig. S12.**
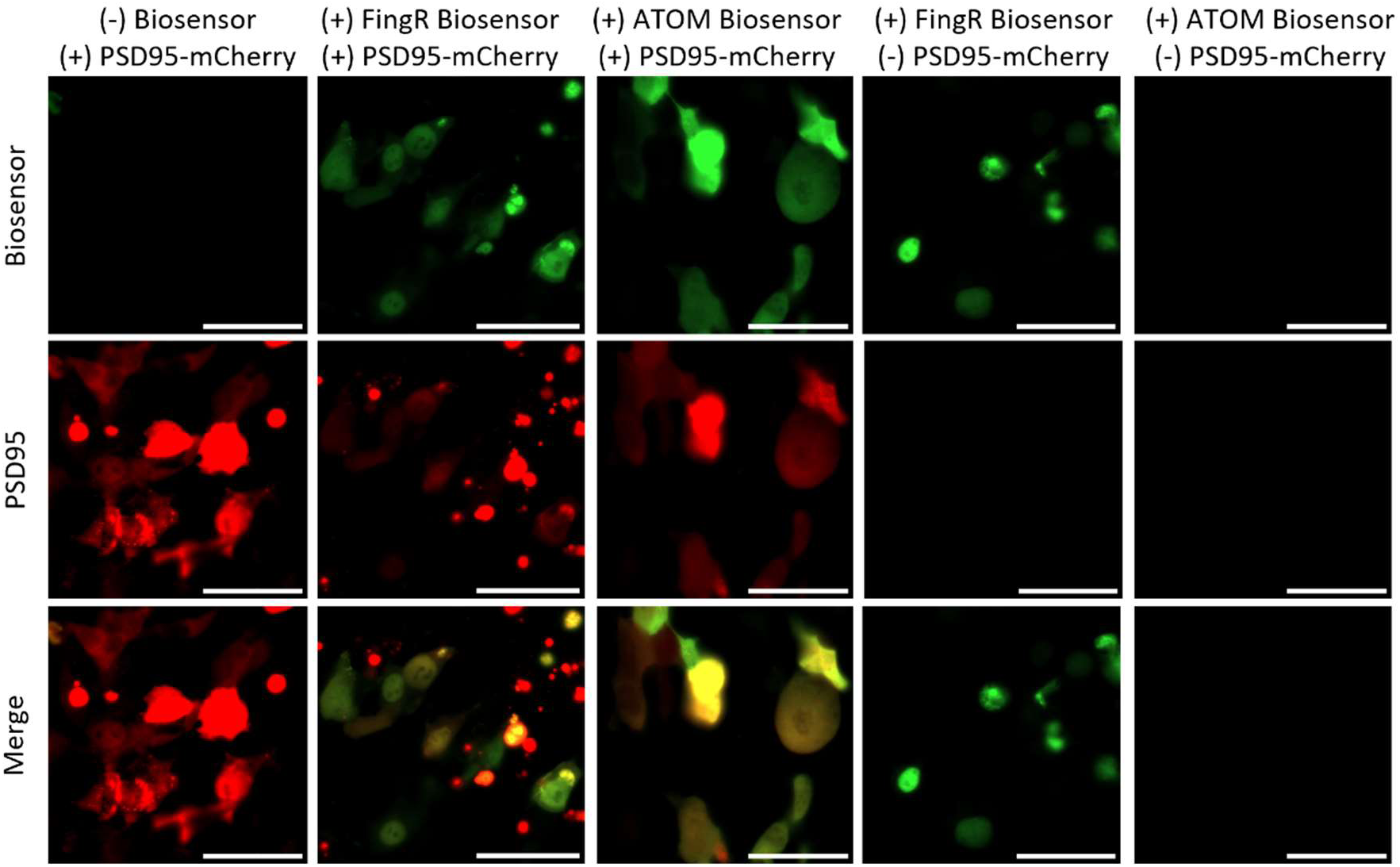
Raw images of the FingR and ATOM biosensors (green channel) co-transfected with PSD95 fused to mCherry (red channel). The FingR biosensor remains mainly in the nucleus without PSD95 but when compared to the no biosensor control, causes PSD95 to form puncta. Surprisingly, the biosensor does not co-localize well with the puncta, suggesting that they might be extracellular. On the other hand, the ATOM biosensor is mainly off in the absence of PSD95 and strongly co-localizes with PSD95 in its presence. The ATOM sensor also retains the diffuse localization of PSD95, as seen in the control.

## Notes

### Competing Interest Statement

The authors have declared no competing interest.

